# Impaired host shutoff is a fitness cost associated with baloxavir marboxil resistance mutations in influenza A virus PA/PA-X nuclease domain

**DOI:** 10.1101/2025.09.22.677688

**Authors:** Jack Case, Denys A. Khaperskyy

**Affiliations:** Department of Microbiology and Immunology, Faculty of Medicine, Dalhousie University, Halifax, NS, Canada

**Keywords:** Baloxavir marboxil, influenza virus, PA-X, polymerase acidic, host shutoff, resistance mutations

## Abstract

The polymerase acidic (PA) protein is a subunit of the trimeric influenza A virus (IAV) RNA-dependent RNA polymerase and the target of the anti-influenza drug baloxavir marboxil (BXM). As with other direct-acting antivirals, treatment with BXM can lead to selection of viruses carrying resistance mutations. If these mutations have negligible fitness costs, resistant viruses can spread widely and render existing treatments obsolete. Multiple BXM resistance mutations in the nuclease domain of PA have been identified, with I38T and I38M amino acid substitutions occurring frequently. These mutations have minimal to no effects on viral polymerase activity, virus replication, or transmission. However, for reasons that are not well understood, viruses with BXM resistance substitutions have not been able to compete with parental wild-type strains. The IAV genome segment encoding PA also encodes the host shutoff nuclease PA-X, which shares the endonuclease domain with PA but has a unique C-terminal domain generated by ribosomal frameshifting during translation. Unlike their effects on PA activity, the effects of BXM or the I38T/M substitutions on PA-X function remain uncharacterized. In our work, for the first time, we directly examine the effects of baloxavir and the I38T/M substitutions on PA-X activity and show that baloxavir inhibits PA-X activity in a dose dependent manner. Most importantly, we also demonstrate that the I38T/M mutations significantly impair the host shutoff activity of PA-X proteins from different IAV strains of H1N1, H3N2, and H5N1 subtypes. Our work reveals that the deleterious effects of I38T/M on PA-X function may represent an important barrier to the spread of BXM-resistant viruses.

**AUTHOR SUMMARY:** A general shut off of the host cell’s ability to produce new proteins, including those that are needed for antiviral defence, is an important feature of the influenza A virus infection. It enables the virus to effectively suppress intrinsic immune responses. Furthermore, influenza viruses continuously evolve and cause seasonal epidemics by escaping adaptive immunity resulting from previous infections or vaccinations. Young children, elderly, and immunocompromised individuals are at increased risk of severe disease and hospitalization following influenza infection. Antiviral drugs are an important option in treating influenza and limiting disease severity. Baloxavir marboxil is a single dose oral anti-influenza drug effective in treatment of influenza A and B. However, viruses frequently evolve resistance to this antiviral, and if resistance spreads, it can render this treatment obsolete. How likely is the emergence of widespread baloxavir resistance remains an open question and in our work we tested if these mutations weaken the virus, and specifically its general host shutoff ability. We discovered that the viral host shutoff is decreased by baloxavir resistance mutations and propose that this could serve as an important barrier to a successful spread of baloxavir-resistant viruses.

## INTRODUCTION

Vaccines to influenza A virus (IAV) were developed decades ago, but due to constant antigenic drift the virus continues to cause seasonal epidemics [1]. In addition to seasonal influenza strains that circulate in humans, the global rise and zoonotic spread of highly pathogenic avian influenza (HPAI) viruses of H5N1 subtype increases the threat of a new pandemic. Large outbreaks in wild birds and poultry in North America, as well as the spread of this virus in cattle in the USA, increase the frequency of human exposure to H5 subtype influenza and have caused numerous infections in people with varied disease severity [2–6]. This underscores the need for better treatment options for those who develop severe respiratory disease, especially in cases where vaccines are ineffective or not yet available. At present there are two classes of anti-influenza drugs widely used in the clinic: neuraminidase inhibitors (oseltamivir, zanamivir, and peramivir) and the polymerase inhibitor baloxavir marboxil (BXM) [7–9]. The latter drug targets the nuclease active site of the viral polymerase acidic (PA) protein located in its N-terminal domain [9]. BXM is effective even as a single oral dose and can be administered in combination with neuraminidase inhibitors [10,11]. Unfortunately, resistance to BXM arises quickly and viruses with resistance mutations represent a continuous threat of rendering this treatment obsolete [9,12–14]. This was the case for adamantanes – once effective antivirals that are no longer used for influenza treatment in the clinic. However, for reasons that are not well understood, viruses with BXM resistance substitutions have not been able to spread widely, despite few documented instances of new infections with BXA-resistant strains [8,13,15,16]. The highest level of resistance is associated with isoleucine 38 to threonine (I38T) substitution in PA (30 to over 100-fold, depending on virus strain and experimental assays), which is also the most frequently detected resistance mutation [8,16]. It is followed by other amino acid changes at this position (e.g. isoleucine to methionine (I38M), phenylalanine (I38F), leucine (I38L), etc.) and substitutions at several other positions in PA (e.g. amino acids 23, 36, 37, 119, 199), however these changes are associated with much lower resistance (1.5 to 9-fold) [17–19]. In the absence of baloxavir, resistance mutations have negligible effects on viral polymerase activity or virus transmission in animal models [16,19]. In *in vitro* cell culture and *in vivo* animal infection models, effects of baloxavir resistance mutations on the efficiency of viral replication and the magnitude of innate antiviral responses vary, but the mutant viruses are usually still able to replicate to high titers [18–20].

The IAV genome segment encoding PA also encodes the host shutoff nuclease PA-X [21]. PA-X is produced from the same messenger RNA (mRNA) as PA via a +1 ribosome frameshift at the phenylalanine-191 codon [22]. This frameshift results in the production of a distinct C-terminal domain called X-ORF, which can be 61 or 41 amino acids in length, depending on the strain [23]. PA-X is the major viral host shutoff factor that mediates cleavage and degradation of host mRNAs and it is conserved in all IAV strains [23]. The endonuclease function of the shared PA/PA-X N-terminal domain is required for PA-X mediated mRNA degradation, while the X-ORF, especially its first 15 amino acids, are important for overall PA-X function by mediating specific protein-protein interactions important for recruiting PA-X towards its target RNAs [24–27]. We have shown previously that PA-X preferentially targets spliced host transcripts that are produced by the cellular RNA polymerase II, and its shutoff activity is augmented by the viral non-structural protein 1 (NS1) – another IAV host shutoff factor [28,29]. NS1 is the major viral innate immune antagonist that interferes with the function of cytoplasmic sensors of viral nucleic acids and disrupts key innate immune signalling pathways [30,31]. It also functions as a general host shutoff factor by interfering with host pre-mRNA processing and nuclear export [32]. The molecular mechanism that links NS1 and PA-X functions is yet to be determined, but several studies have shown that the magnitudes of PA-X and NS1 host shutoff must be balanced for optimal viral replication [33–35].

Because the N-terminal nuclease domain is shared between PA and PA-X, any baloxavir resistance substitutions that emerge would be present in both proteins. While the effects of these mutations on PA function have been reported, no studies have directly examined the effects of baloxavir resistance mutations on PA-X. In this work, we tested the effects of I38T and I38M substitutions in the PA gene segment on PA-X shutoff activity using transfection-based reporter shutoff assays and cell culture infection models with recombinant mutant viruses. We selected these two mutations because they occur most frequently and confer the highest levels of baloxavir resistance. Our work reveals that both mutations impair PA-X host shutoff activity, with the I38T substitution having the most profound effect, completely inactivating PA-X of the laboratory adapted H1N1 strain. The baseline activity of PA-X and the negative effect of baloxavir resistance mutations on PA-X shutoff varied between different virus strains. However, I38T and I38M decreased PA-X activity in all PA-X variants tested, indicating that the impaired PA-X mediated host shutoff may represent an important barrier to the establishment and spread of the BXM-resistant IAVs.

## RESULTS

### Host shutoff endonuclease PA-X is inhibited by baloxavir

The endonuclease active site targeted by baloxavir is shared between PA and PA-X proteins (Fig. 1A). Therefore, we tested if baloxavir acid (BXA), a cell-permeable active metabolite of the baloxavir marboxil pro-drug, inhibits activity of the PA-X protein in our established transfection-based shutoff assay. We co-transfected HEK 293A cells with the c-terminally myc-tagged PA-X expression construct from the A/PuertoRico/8/34(H1N1) strain (PR8) or an empty vector control with the EGFP reporter plasmid. At 4 h post-transfection, we treated cells with increasing concentrations of BXA and measured GFP signal at 24 h post-transfection (Fig. 1B,C). As expected, GFP signal in PA-X transfected cells was strongly decreased compared to control. This shutoff of reporter expression was reversed by treatment with BXA in a concentration-dependent manner, with 200 nM BXA completely blocking the PA-X mediated inhibition of EGFP expression (Fig. 1B,C). Treatment of cells with BXA did not interfere with reporter expression in control transfections (Fig. 1B,C) and its effect on PA-X host shutoff was not due to decreased PA-X expression (Fig. 1D). Instead, PA-X protein levels were increased with BXA treatment due to a relief of the PA-X mediated shutoff of its

**Figure 1.**
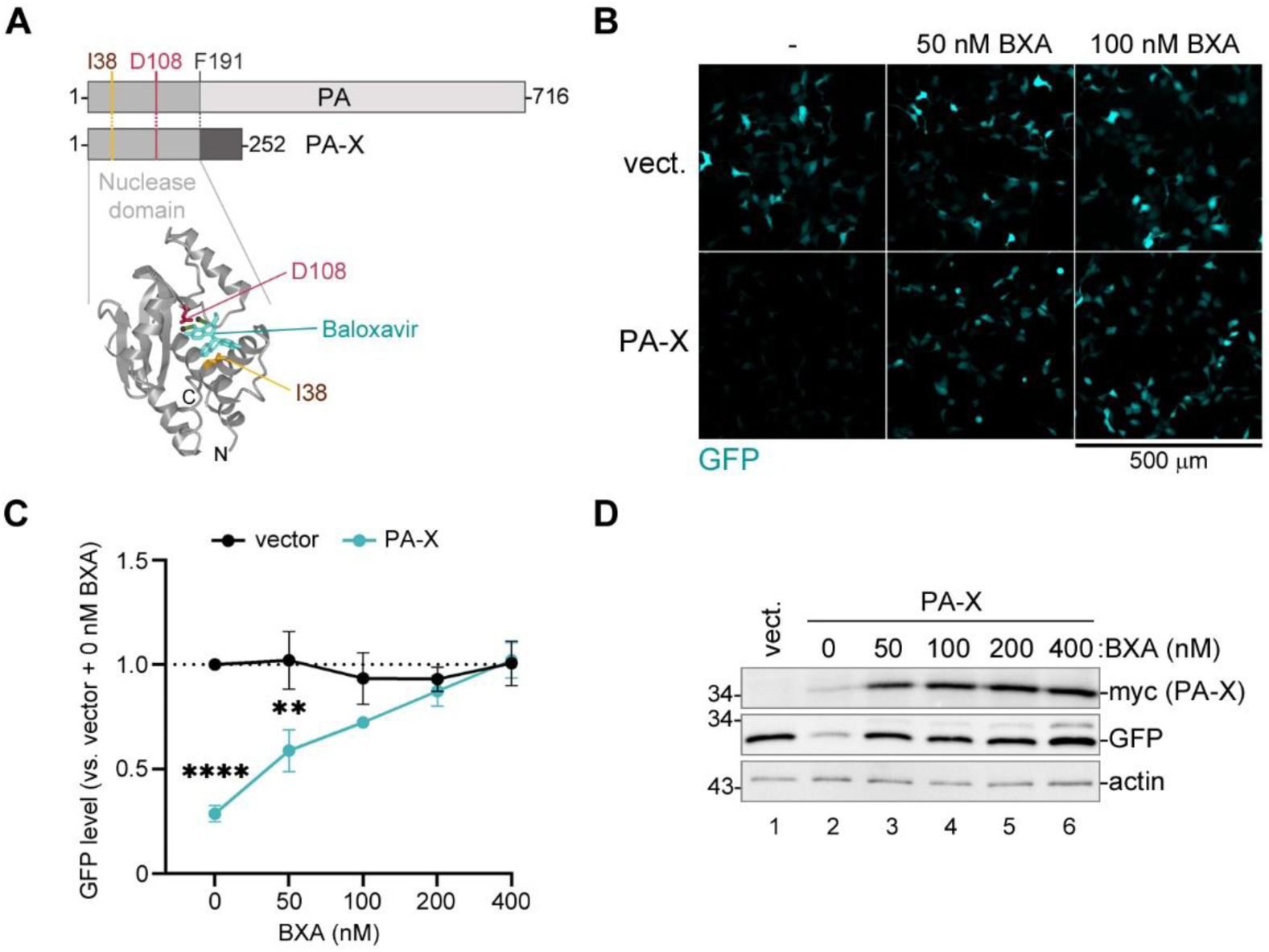
Baloxavir inhibits PA-X shutoff activity. (A) PA and PA-X share the nuclease domain and any amino acid substitutions in the active site. Top: linear diagram of PA and PA-X proteins. Position of the F191 amino acid that marks the end of the shared sequence between PA and PA-X is indicated. Bottom: ribbon diagram of the nuclease domain structure with baloxavir acid (teal) bound in the active site (PDB ID: 6FS6) [12]. Side chains and positions of the conserved amino acids in the active site D108 (essential for nuclease activity) and I38 (site of baloxavir resistance mutations) are highlighted in red and brown, respectively. (B-D) 293A cells were co-transfected with the EGFP reporter plasmid and either the myc-tagged PA-X expression construct derived from the PR8 strain (PA-X) or the empty vector control (vect.). At 4 h post-transfection, cells were treated with the indicated concentrations of balaxovir acid (BXA) or vehicle control (-) and analysed at 24 h post-transfection. (B) Representative fluorescence microscopy images showing reporter expression. Scale bar = 500 µm. (C) The relative signal intensity in the GFP channel was quantified from images represented in panel B (N = 3, see materials and methods section for details). One-way ANOVA and Dunnett multiple comparisons tests were used to determine statistical significance (****P-value <0.0001; **P-value < 0.01). Error bars = standard deviation. (D) Whole cell lysates were collected from transfected cells treated as indicated and the expression of EGFP and PA-X was visualised by western blotting with antibodies to GFP and the myc tag, respectively. Actin was used as a loading control. own expression (Fig. 1D). Thus, our reporter shutoff assay shows that the PA-X activity is inhibited by baloxavir.

### Isoleucine 38 substitutions that confer baloxavir resistance impair PA-X shutoff activity

To test the effects of mutations that confer the highest levels of baloxavir resistance to the viral RdRp on the shutoff activity of PA-X, we introduced isoleucine 38 to threonine (38T) or methionine (38M) substitutions into our PR8 PA-X expression construct and used them in our reporter shutoff assay. As an additional control, we used catalytically inactive aspartate 108 to alanine (108A) PA-X mutant (Fig. 2A-C). In this assay, all mutations strongly decreased PA-X host shutoff, with the 38T substitution inactivating PA-X to a similar degree as the 108A substitution, while the 38M PA-X mutant retained some host shutoff activity (Fig. 2A,B). The mutations also caused increased PA-X protein accumulation compared to the wild-type (WT) due to the relief of the shutoff of its own expression (Fig. 2C). Together, these results show that the 38T mutation, which confers the highest degree of baloxavir resistance, also causes the strongest impairment to PA-X function in our system. To confirm that the 38T and 38M substitutions result in a high degree of baloxavir resistance without strong effects on the function of the PR8 RdRp, we performed a luciferase replicon assay using PB2, PB1, and NP proteins from this strain in combination with the C-terminally myc-tagged WT, 38T, or 38M PA. Consistent with previous studies, we observed comparable activities between the WT and mutant PA RdRp complexes in the absence of BXA treatment (Fig. 2D). The WT PR8 RdRp activity sharply decreased with increasing concentrations of BXA (IC50 = 3.85±1.53 nM based on a linear regression analysis). The 38M PA substitution conferred 9-fold resistance in our assay (IC50 = 34.51±3.34 nM), and 38T mutation conferred 26-fold resistance (IC50 = 99.38±6.29 nM) (Fig. 2D). These results demonstrate that 38T and 38M substitutions in the nuclease domain that impair PA-X shutoff activity do not significantly impair RdRp activity and confer strong BXA resistance, as expected based on previous studies [12,18,19].

**Figure 2.**
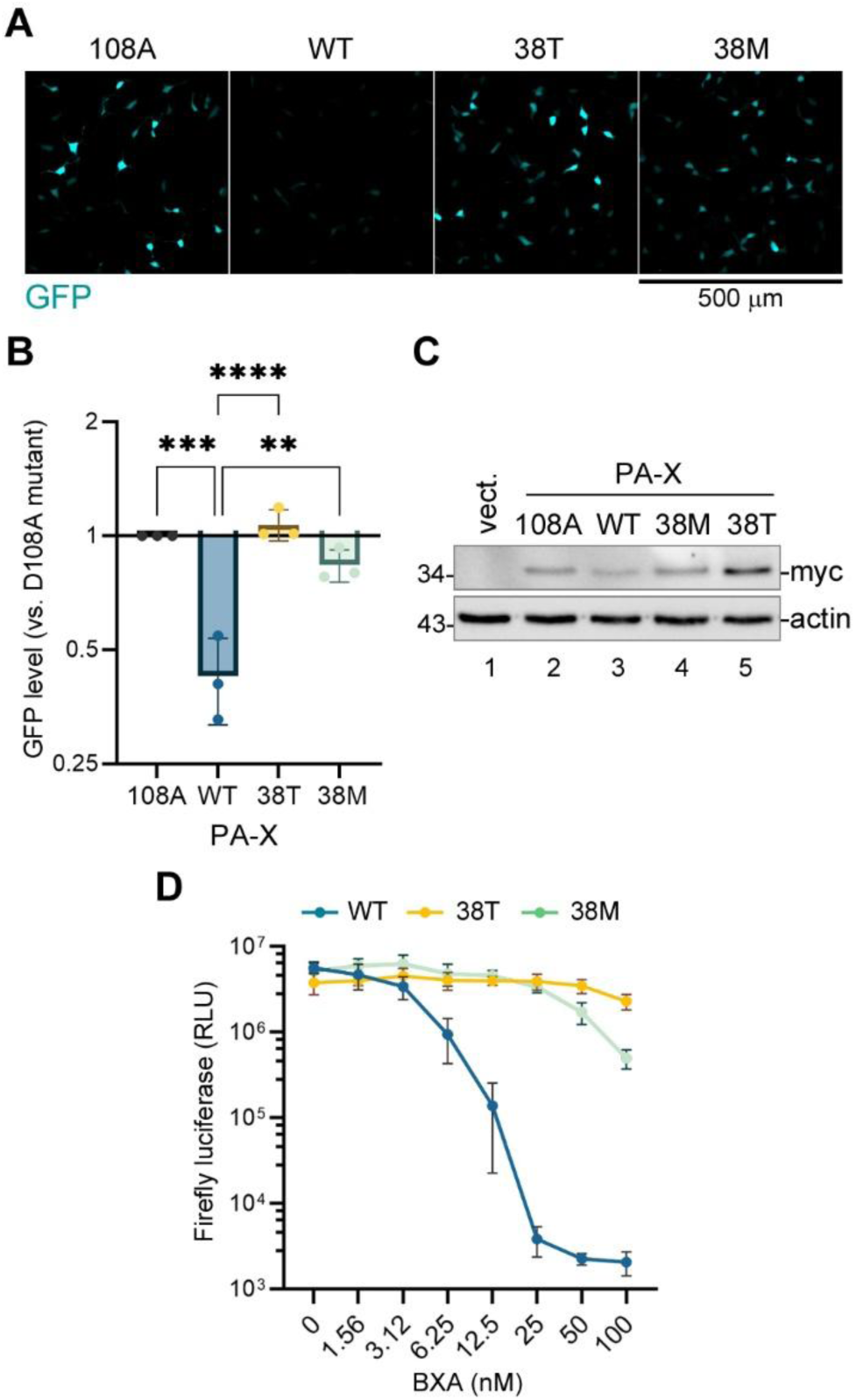
Baloxavir resistance mutations impair PA-X shutoff activity. (A-C) 293A cells were co-transfected with the EGFP reporter plasmid and the indicated myc-tagged wild-type (WT) or mutant PA-X expression construct derived from the PR8 strain and analysed at 24 h post-transfection. (A) Representative fluorescence microscopy images showing reporter expression. Scale bar = 500 µm. (B) The relative signal intensity in the GFP channel was quantified from images represented in panel A (N = 3, see materials and methods section for details). One-way ANOVA and Dunnett multiple comparisons tests were used to determine statistical significance (****P-value <0.0001; ***P-value < 0.001; **P-value < 0.01). Error bars = standard deviation. (C) Whole cell lysates were collected from cells transfected with the indicated PA-X expression constructs or with empty vector (vect.) and the expression of PA-X was visualised by western blotting with antibody to myc tag (myc). Actin was used as a loading control. (D) 293A cells were co-transfected with the firefly luciferase influenza replicon vector, the PR8 NP, PB2, PB1 expression constructs, and the indicated wild-type (WT) or mutant myc-tagged PR8 PA expression plasmids. At 4 h post-transfection, cells were treated with the indicated concentrations of balaxovir acid (BXA) and the luciferase activity was measured 24 h post-transfection. RLU = relative luminescence units.

### Baloxavir resistance mutations confer phenotypes similar to a frame shift site mutation that disrupts PA-X production

Previously, we generated the PA-X deficient mutant virus PR8-PA(fs) by introducing two synonymous nucleotide substitutions in the PA-X frame shift site [24] and characterized its replication and host shutoff phenotypes in an A549 cell culture infection model in comparison to a WT PR8 virus [26,28,29]. One of the striking phenotypes associated with PA-X host shutoff is the nuclear relocalization of the cytoplasmic poly(A) binding protein (PABPC1). This phenotype correlates with the magnitude of the host mRNA depletion and is strongly attenuated in PR8-PA(fs) virus infected cells compared to WT [24,29]. To examine the effects of baloxavir resistance mutations on viral host shutoff and compare them to the PA(fs) mutation, we produced recombinant PR8 viruses with PA(38T) and PA(38M) substitutions. Then, we infected A549 cells with the WT PR8, PR8-PA(fs), PR8-PA(38T), and PR8-PA(38M) viruses at a multiplicity of infection (MOI) of 1 and analyzed virus and mock infected cells at 21 h post-infection (hpi). Using immunofluorescence microscopy, we measured the nuclear PABPC1 accumulation and observed that approximately half of the WT PR8 virus infected cells had nuclear PABPC1 at 21 hpi (Fig. 3A,B). The fraction of infected cells with nuclear PABPC1 decreased significantly by PA(fs) mutation as well as PA(38T) and PA(38M) mutation, with the PA(38T) mutation having the strongest effect (Fig. 3B). This indicates that the magnitude of host shutoff is significantly decreased by baloxavir resistance mutations, and this decrease is comparable or even exceeds that caused by PA-X deficiency. Importantly, the decreased host shutoff was not caused by lower viral protein accumulation in mutant virus infected cells (Fig. 3C). To further confirm that the host shutoff is affected, we compared levels of the cellular *ACTB* and *G6PD* transcripts in mock, WT, and mutant virus infected cells using RT-qPCR (Fig. 3D,E). We have previously shown that these mRNAs are downregulated by PA-X in virus-infected cells [28] and, in this experiment, their depletion in cells infected with either PA(fs) or PA(38T) mutant viruses was strongly attenuated compared to the WT virus-infected cells (Fig. 3D,E). Taken together, these results confirm our findings using reporter co-transfection assays that baloxavir resistance mutations at isoleucine 38 in the PA endonuclease domain significantly impair viral host shutoff.

**Figure 3.**
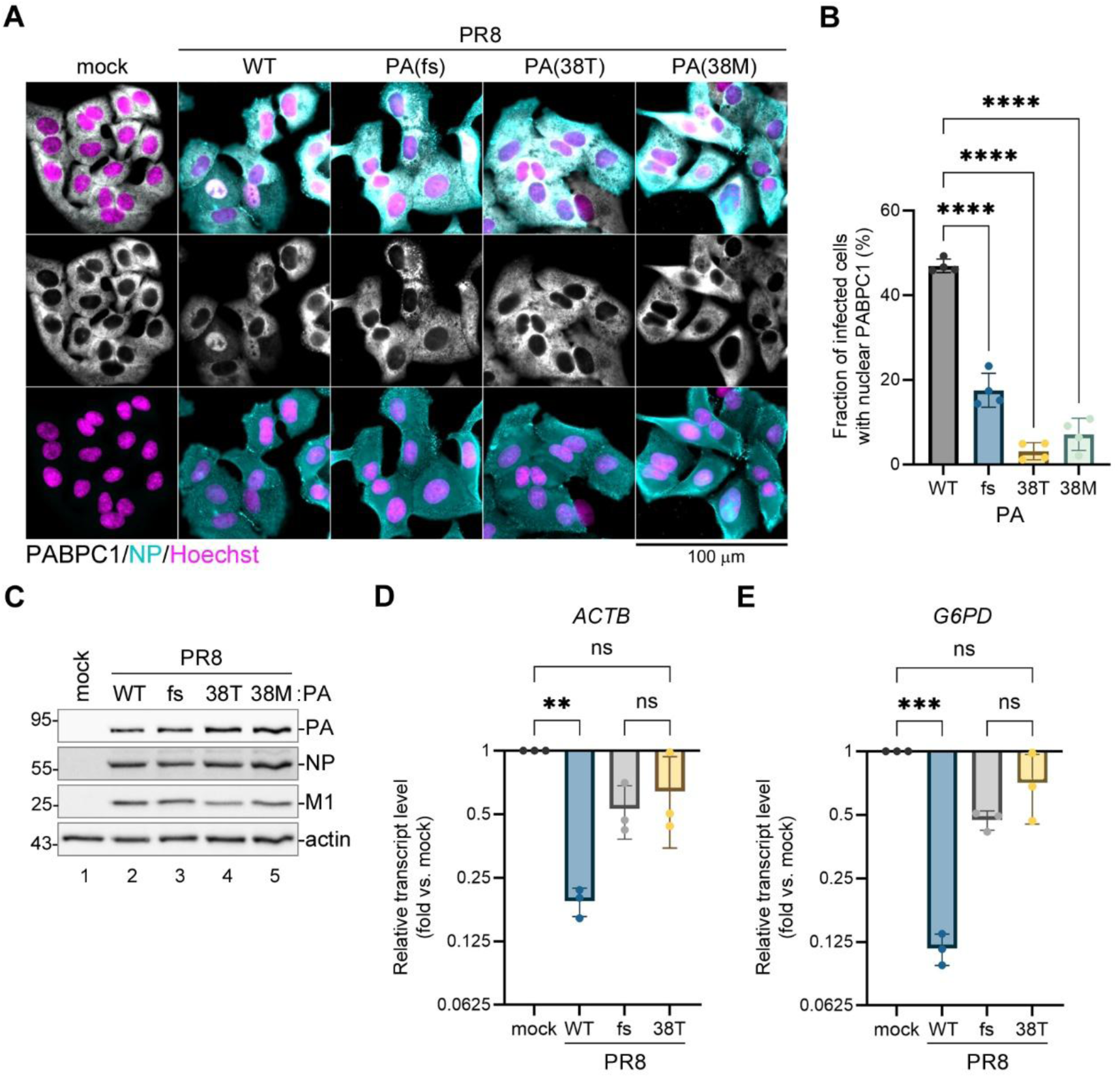
Baloxavir resistance mutations impair host shutoff by a laboratory-adapted influenza virus strain. A549 cells were infected with the reverse-genetic derived PR8 wild-type (WT) or the indicated PA mutant influenza viruses at a multiplicity of infection (MOI) of 1.0 or mock infected and analyzed at 21 h post-infection. (A) Immunofluorescence microscopy images showing subcellular distribution of the cytoplasmic poly(A) binding protein (PABPC1, white). Infected cells are immunostained for NP expression (teal), nuclei are stained with Hoechst dye (purple). Scale bar = 100 µm. (B) Fraction of infected cells with nuclear PABPC1 accumulation was quantified from images presented in panel A (N = 4, see materials and methods section for details). (C) Accumulation of viral proteins in infected cells was visualized by western blotting using the indicated antibodies. Actin was used as a loading control. (D,E) Relative levels of host *ACTB* (D) and *G6PD* (E) transcripts in mock and virus-infected cells was determined using RT-qPCR (N = 3). In B, D, and E, one-way ANOVA and Dunnett multiple comparisons tests were used to determine statistical significance (****P-value <0.0001; ***P-value <0.001; **P-value < 0.01; ns: non-significant). Error bars = standard deviation.

### Substitutions that confer baloxavir resistance attenuate host shutoff by the 2009 pandemic H1N1 virus

PA-X activity and the magnitude of host shutoff vary significantly between influenza virus strains. To confirm that the baloxavir resistance mutations I38T/M affect PA-X activity in other H1N1 strains, we cloned PA-X from the A/California/7/09(H1N1) (CA07) 2009 pandemic virus strain, introduced 38T and 38M substitutions, and conducted EGFP reporter shutoff assays (Fig. 4A,B). Both baloxavir resistance mutations significantly attenuated CA07 PA-X activity compared to WT PA-X (Fig. 4B) and, as a consequence, these mutant PA-X proteins were expressed at higher levels than the WT CA07 PA-X upon transient transfection (Fig. 4A). However, the magnitude of the reporter shutoff by the CA07 PA-X was approximately 2-fold higher than by the PR8 PA-X, and both the 38T and 38M mutant CA07 PA-X proteins significantly downregulated EGFP reporter expression (Fig. 4B).

**Figure 4.**
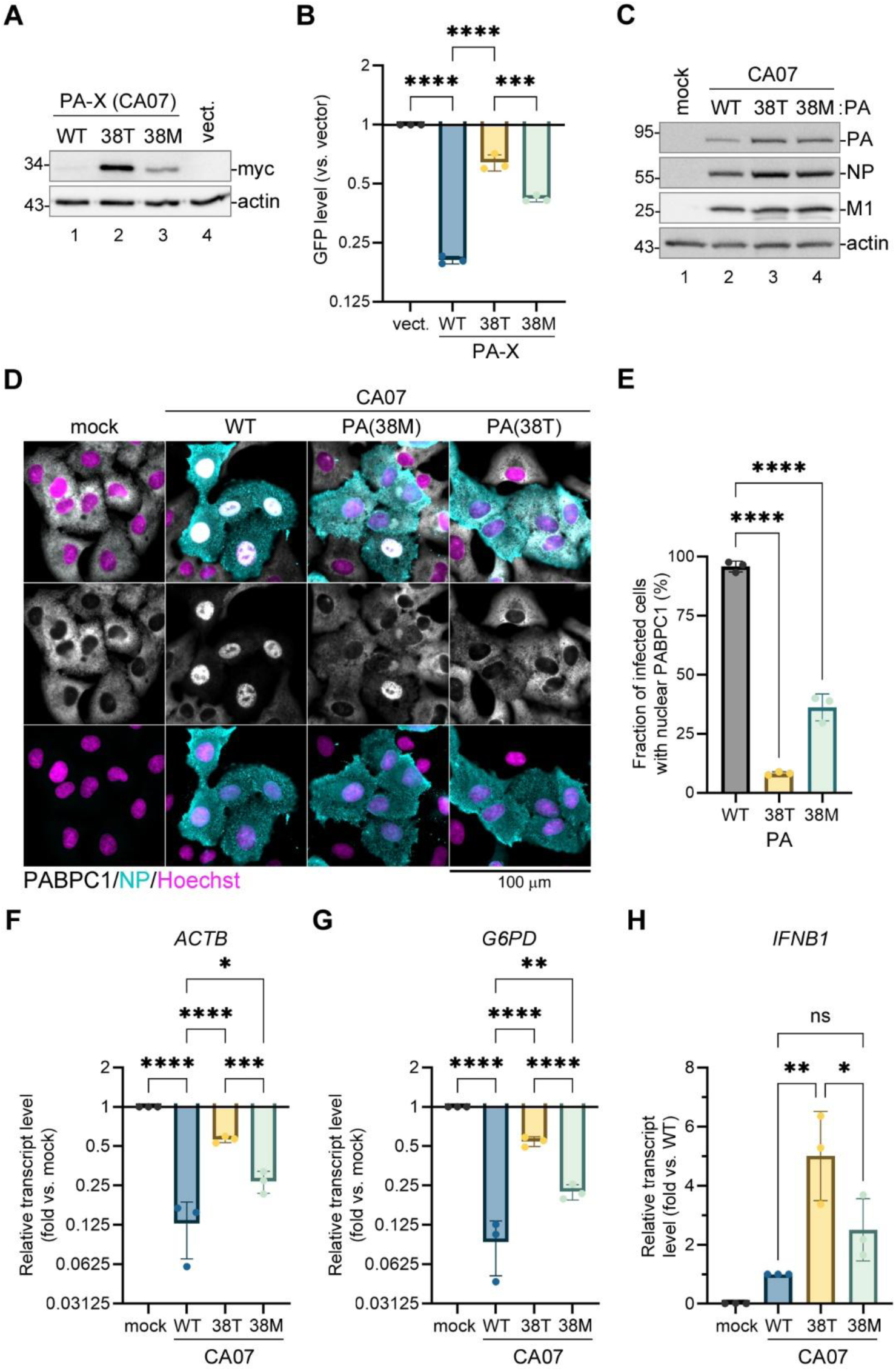
Baloxavir resistance mutations significantly attenuate host shutoff by the 2009 pH1N1 strain. (A-B) 293A cells were co-transfected with the EGFP reporter plasmid and the indicated myc-tagged wild-type (WT) or mutant PA-X expression construct derived from the CA07 strain and analysed at 24 h post-transfection. (A) Whole cell lysates were collected from cells transfected with the indicated PA-X expression constructs or with empty vector (vect.) and the expression of PA-X was visualised by western blotting with antibody to myc tag (myc). Actin was used as a loading control. (B) The relative signal intensity in the GFP channel was quantified from fluorescence microscopy images (N = 3, see materials and methods section for details). (C-H) A549 cells were infected with the reverse-genetic derived CA07 wild-type (WT) or the indicated PA mutant influenza viruses at a multiplicity of infection (MOI) of 1.0 or mock infected and analyzed at 21 h post-infection. (C) Accumulation of viral proteins in infected cells was visualized by western blotting using the indicated antibodies. Actin was used as a loading control. (D) Immunofluorescence microscopy images showing subcellular distribution of the cytoplasmic poly(A) binding protein (PABPC1, white). Infected cells are immunostained for NP expression (teal), nuclei are stained with Hoechst dye (purple). Scale bar = 100 µm. (E) Fraction of infected cells with nuclear PABPC1 accumulation was quantified from images presented in panel D (N = 3, see materials and methods section for details). (F-H) Relative levels of host ACTB (F), G6PD (G), and IFNB1 (H) transcripts in mock and virus-infected cells was determined using RT-qPCR (N = 3). In B and E-H, one-way ANOVA and Dunnett multiple comparisons tests were used to determine statistical significance (****P-value <0.0001; ***P-value < 0.001; **P-value < 0.01; *P-value < 0.05; ns: non-significant). Error bars = standard deviation.

Of the two mutations, 38M had a significantly weaker effect on PA-X activity than the 38T substitution, and the activity of CA07 PA-X(38M) was comparable to the activity of the WT PR8 PA-X in this assay (compare Fig. 4B to 2B). This indicates that the CA07 virus with baloxavir resistance substitutions may still be able to induce a significant level of host shutoff in infected cells. To test this, we used reverse genetics to produce CA07-PA(38T) and CA07-PA(38M) viruses and compare their host shutoff phenotypes to the WT CA07 virus in a cell culture infection model (Fig. 4C-H). Similar to the PR8 infection, we did not observe a decrease in viral protein accumulation but instead detected slightly higher levels of PA and NP proteins in CA07-PA(38T) and CA07-PA(38M) infected cells compared to WT virus-infected cells (Fig. 4C). Of note, in several previous studies, increased PA expression was also associated with mutations that disrupt PA-X production [29,36,37]. Consistent with a high magnitude of host shutoff by the CA07 virus, nearly all WT virus-infected cells had strong nuclear PABPC1 accumulation at 21 hpi (Fig. 4D,E). This phenotype was significantly attenuated in cells infected with CA07-PA(38M) mutant virus and even more strongly impaired in cells infected with CA07-PA(38T) virus, reflecting the different degree of the negative effect on PA-X activity. Similarly, the depletion of host *ACTB* and G6PD transcripts was significantly affected by baloxavir resistance mutations, with 38T having a stronger negative effect compared to 38M substitution (Fig. 4F,G). To test if lower host shutoff results in higher induction of antiviral responses in cells infected with viruses with these baloxavir resistance mutations, we measured levels of interferon-β transcript (*IFNB1*) in infected cells. We detected significantly higher induction of *IFNB1* (∼5-fold) in cells infected with the CA07-PA(38T) virus compared to the WT, and on average a 2-fold increased *IFNB1* transcript levels in CA07-PA(38M) infected cells which did not reach statistical significance (Fig. 4H). Previously, increased interferon induction was shown in a study analyzing viruses with baloxavir resistance mutations in cell culture [18], and our results are in agreement with these findings. In our model, we also observe that, although CA07 PA-X activity and the magnitude of host shutoff are decreased by the 38T and 38M baloxavir resistance substitutions, mutant viruses are still inducing a measurable degree of host shutoff, and the two mutations vary significantly in their fitness costs.

### Significant shutoff activity is preserved in H3N2 and H5N1 strain-derived PA-X proteins with 38T and 38M substitutions

Having observed differences in the magnitude of host shutoff and the effects of baloxavir resistance substitutions between the laboratory-adapted and pandemic H1N1 strains, we extended our analyses to PA-X proteins from the H3N2 and H5N1 viruses: A/Switzerland/9715293/13(H3N2) (SW13) and A/Bovine/texas/24-029328-01/24(H5N1) (TX24). Although the PA-X amino acid sequences from these strains are over 90% identical to PR8 and CA07 PA-X proteins, they possess several key changes in the N-terminal nuclease domain that can affect their activity (S1 Fig.). We cloned the PA-X sequences from these strains in the same C-terminally myc-tagged expression vector that we used for the PR8 and CA07 PA-X overexpression and conducted EGFP reporter shutoff assays (Fig. 5A,B). Based on our fluorescence microscopy images, the SW13 PA-X had a very potent shutoff activity, which was decreased by the 38T and, to a lesser extent, by the 38M substitutions (Fig. 5A). However, when we quantified the GFP fluorescence signal using our image processing pipeline, the changes due to 38T and 38M substitutions that were consistently observed visually did not appear significant (Fig. 5B). This was likely due to a sharp decrease in the dynamic range of our analyses at the lower fluorescence intensities. To mitigate this limitation, we repeated our co-transfection shutoff assay with an intron-containing firefly luciferase reporter used in our previous study [26]. For this analysis, we also included the CA07 PA-X constructs as a comparative control and a PA-X constructs from the recent human H5N1 isolate A/BC/PHL-2032/24(H5N1) (BC24) that has a different origin PA segment than the TX24 PA [2] (Fig. 5C-E). This analysis revealed significant attenuation of the host shutoff due to the 38T and 38M substitutions in all PA-X constructs except for SW13 PA-X(38M) which was not significant due to a higher replicate variability (Fig. 5C). Interestingly, SW13 PA-X caused the strongest luciferase reporter downregulation, and CA07 PA-X was the least potent. Both of the H5N1 PA-X proteins had strong shutoff activity that was significantly reduced by the baloxavir resistance mutations, but this reduced activity was still comparable to the WT CA07 PA-X (Fig. 5C). As with the

**Figure 5.**
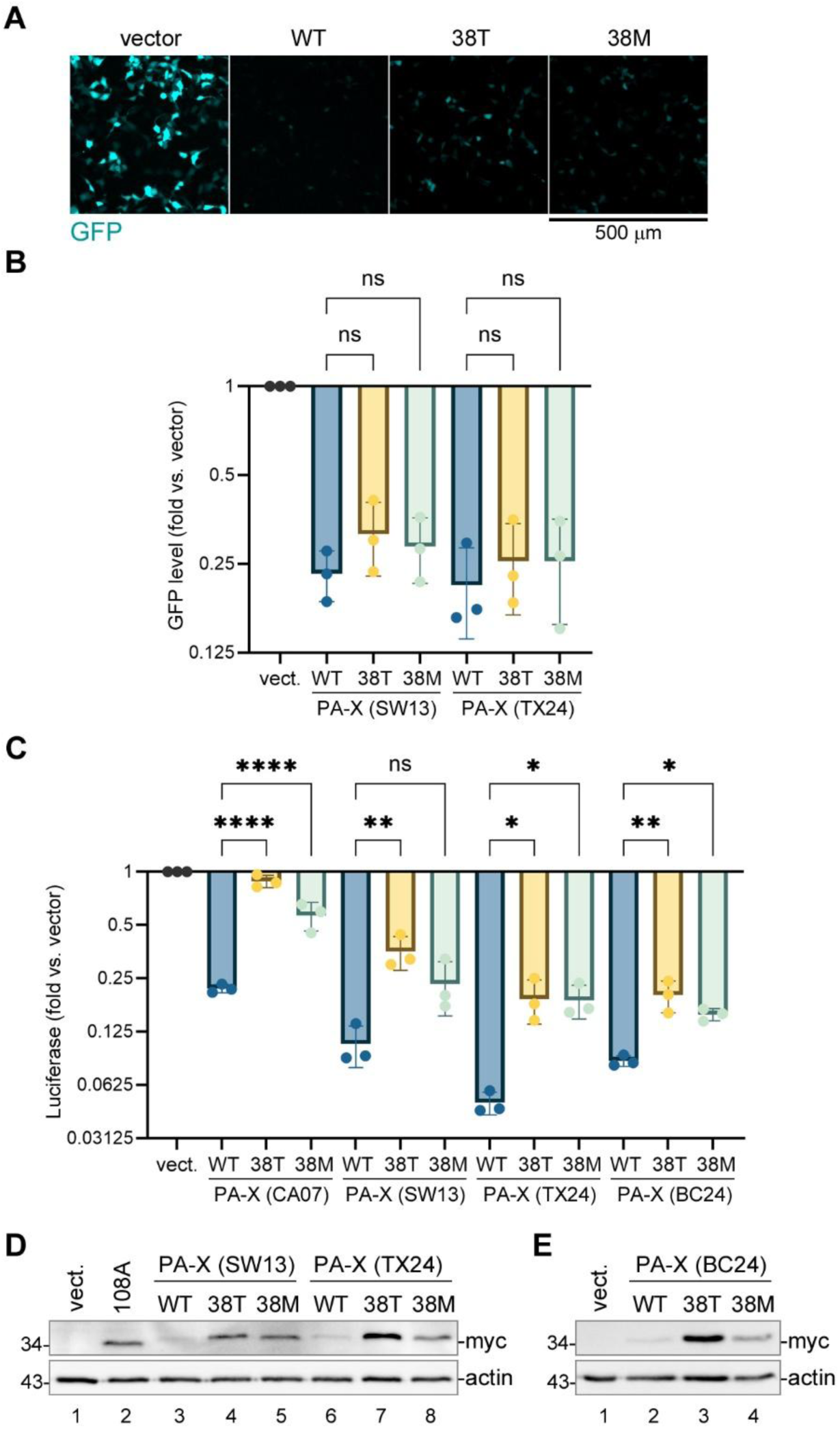
PA-X proteins from H3N2 and H5N1 influenza virus strains have higher shutoff activity compared to H1N1-derived PA-X. (A,B) 293A cells were co-transfected with the EGFP reporter plasmid and the indicated myc-tagged wild-type (WT) or mutant PA-X expression constructs derived from the SW13 (H3N2) or TX24 (H5N1) strains and analysed at 24 h post-transfection. (A) Representative fluorescence microscopy images showing reporter expression in cells co-transfected with the indicated WT or mutant SW13 PA-X constructs. Scale bar = 500 µm. (B) The relative signal intensity in the GFP channel was quantified from fluorescence microscopy images (N = 3, see materials and methods section for details). (C-E) 293A cells were co-transfected with the firefly luciferase reporter plasmid and the indicated myc-tagged wild-type (WT) or mutant PA-X expression constructs derived from the CA07 (H1N1), SW13 (H3N2), TX24 (H5N1), or BC24 (H5N1) strains and analysed at 24 h post-transfection. (C) The relative firefly luciferase activity was measured (N = 3). (D,E) Whole cell lysates were collected from cells transfected with the indicated PA-X expression constructs, empty vector (vect.), or catalytically inactive D108A mutant PR8 PA-X (108A) and the expression of PA-X was visualised by western blotting with antibody to myc tag (myc). Actin was used as a loading control. In B and C, one-way ANOVA and Dunnett multiple comparisons tests were used to determine statistical significance (****P-value <0.0001; **P-value < 0.01; *P-value < 0.05; ns: non-significant). Error bars = standard deviation.

PR8 and CA07 PA-X proteins analysed before (Fig. 2C, 4A), a decrease in shutoff activity resulted in higher accumulation of the 38T and, to a lesser extent, of the 38M mutant PA-X proteins compared to the WT PA-X (Fig 5D,E). This makes a direct quantitative comparison of the host mRNA cleavage and degradation activities between the WT and mutant PA-X proteins difficult, as a lower activity can be partially compensated by an increased expression and vice versa. Nevertheless, based on our reporter assay, among different baloxavir resistance mutants, the BC24 PA-X(38M) protein retained the highest shutoff activity.

### Baloxavir resistance mutations in the PA segment preserve H5N1 virus RdRp activity but significantly attenuate PA-X host shutoff

Because of a significant retention of the reporter shutoff activity by the ectopically expressed BC24 PA-X with the 38T or 38M baloxavir resistance mutations, we constructed a reconstituted BC24 minireplicon to examine the effects of these mutations on PA activity in the RdRp complex and on the activity of PA-X produced in this system via a natural low-frequency +1 frame shift. Using the minireplicon mitigates potential biosafety concerns of introducing baloxavir resistance mutations into infectious H5N1 highly pathogenic influenza virus while better mimicking the mechanism of PA-X production and function in infected cells compared to PA-X overexpression. First, to confirm that the 38T and 38M substitutions confer baloxavir resistance, we performed a luciferase replicon assay in the presence of varying concentrations of BXA using BC24 PB2, PB1, and NP proteins in combination with the c-terminally myc-tagged WT, 38T, or 38M BC24 PA (Fig. 6A). Similar to the PR8 replicon (Fig. 2D), the WT BC24 replicon activity was highly sensitive to BXA treatment (EC50 = 1.67±0.09 nM). The 38M substitution conferred ∼9-fold resistance (EC50 = 15.64±5.26 nM), and the 38T substitution conferred nearly 60-fold resistance (EC50 = 98.91±20.54 nM). In addition, levels of the firefly replicon reporter (Fig. 6A) and the PA proteins (Fig. 6B) in the absence of BXA treatment were higher in cells co-transfected with the 38T or 38M mutant PA constructs compared to the WT. We speculate that this effect is due to a reduced PA-X shutoff activity associated with 38T and 38M substitutions that permits better expression of the replicon components by transfected cells, similar to the effects of these substitutions on the ectopic expression of PA-X (Fig. 5D,E). Next, we substituted the firefly replicon reporter with the EGFP replicon reporter and performed immunofluorescence microscopy analysis of subcellular PABPC1 distribution in transfected cells (Fig. 6C,D). In this assay, GFP fluorescence marks cells with a successful reconstitution of the viral RdRp function, and, in our system, over 40% of the GFP-positive cells had nuclear PABPC1 relocalization, indicative of a robust PA-X mediated host shutoff (Fig. 6D). This nuclear PABPC1 redistribution was significantly decreased by the 38M substitution and strongly inhibited by the 38T substitution in the PA segment (Fig. 6C,D). Notably, many GFP-positive cells co-transfected with the 38T mutant PA displayed bright cytoplasmic PABPC1 foci (Fig. 6C). These foci are most likely stress granules – condensates of untranslated mRNAs, translation factors, and select RNA binding proteins that form as a result of stress-induced translation arrest [38,39]. In our previous work, we demonstrated that the PA-X mediated host shutoff is important in preventing stress granule formation in influenza virus-infected cells [24], and the appearance of these cytoplasmic PABPC1 foci further indicates decreased PA-X activity caused by the 38T substitution.

**Figure 6.**
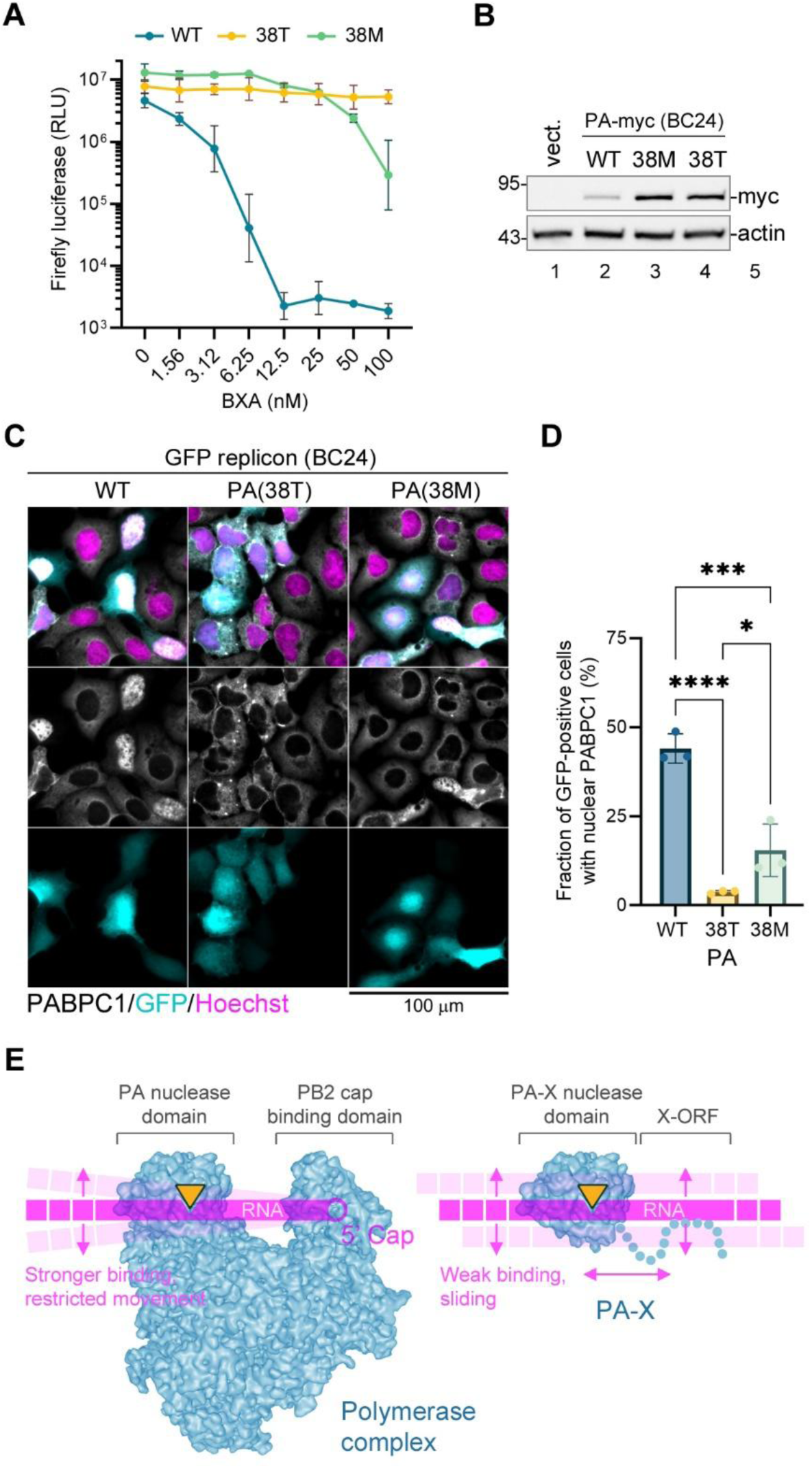
Baloxavir resistance mutations have stronger negative effects on PA-X function compared to PA function. (A,B) 293A cells were co-transfected with the firefly luciferase influenza replicon vector, the BC24 NP, PB2, PB1 expression constructs, and the indicated wild-type (WT) or mutant myc-tagged BC24 PA expression plasmids. At 4 h post-transfection, cells were treated with the indicated concentrations of balaxovir acid (BXA) and analyzed 24 h post-transfection. (A) The firefly luciferase activity was measured (N = 3). RLU = relative luminescence units. (B) Expression of PA in transfected cells was visualized by western blotting using anti-myc (myc) antibody. Actin was used as a loading control. (C-D). 293A cells were co-transfected with the EGFP influenza replicon vector, the BC24 NP, PB2, PB1 expression constructs, and the indicated wild-type (WT) or mutant myc-tagged BC24 PA expression plasmids and analyzed at 24 h post-transfection. (C) Immunofluorescence microscopy images showing subcellular distribution of the cytoplasmic poly(A) binding protein (PABPC1, white). Cells with active reconstituted replicon are visualized by GFP fluorescence (teal), nuclei are stained with Hoechst dye (purple). Scale bar = 100 µm. (D) Fraction of GFP-positive cells with nuclear PABPC1 accumulation was quantified from images presented in panel C (N = 3, see materials and methods section for details). One-way ANOVA and Dunnett multiple comparisons tests were used to determine statistical significance (****P-value <0.0001; ***P-value < 0.001; *P-value < 0.05). Error bars = standard deviation. (E) A model diagram for the differential effects of baloxavir resistance mutations on PA vs. PA-X function. Left: RNA polymerase cap snatching function involves coordinated action of the PB2 cap binding and the PA nuclease domains. Capture of the pre-mRNA 5’ end cap by the PB2 cap binding domain restricts movement of the RNA and orients it towards the PA endonuclease domain for optimal cleavage. Right: transient interaction of PA-X with RNA requires high nuclease activity for efficient cleavage. Structures for the trimeric RNA polymerase complex and the PA endonuclease domain are derived from PDB ID 7NHX [54]. Nuclease active site is represented by an orange triangle.

## DISCUSSION

Our data shows that the acquisition of high baloxavir resistance through adaptive substitutions of threonine or methionine for isoleucine 38 in the PA active site leads to impaired PA-X activity. Although we and others have shown that the PA-X deficient mutant viruses replicate to high titers in cell culture and mouse infection models, they elicit stronger innate and adaptive antiviral responses [21,28,40,41]. The PA-X open reading frame is preserved in all IAV strains and subtypes, indicating that this factor is important for the viral evasion of host immunity and overall circulation of IAV in nature [23]. Accordingly, impairment of PA-X mediated host shutoff must represent a barrier to the successful spread of baloxavir-resistant IAVs.

Our results also show that the shutoff activity of PA-X from different IAV strains varies significantly. In a laboratory adapted PR8 strain PA-X activity is low and it is completely inactivated by the 38T substitution. By contrast, we observed strong shutoff activities of the PA-X proteins from the H3N2 strain and from the highly pathogenic avian-origin H5N1 viruses that was not fully inhibited by the 38T mutation. The PA-X proteins of H1N1 viruses encoded by the PA segments of avian origin, such as CA07 [42], were previously shown to have higher shutoff activity compared to PA-X proteins of their descendant human-adapted seasonal H1N1 strains [33]. These differences were attributed to amino acid changes at amino acid positions 100, 204, 221, and 229 [33]. Of those, valine-100 in the N-terminal nuclease domain is present in CA07 and the H5N1 strains TX24 and BC24. This residue is an alanine in PR8 and H3N2 PA-X proteins, and an isoleucine in the currently circulating seasonal H1N1 strains. Given the high PA-X shutoff activity of SW13 (H3N2) and higher activity of H5N1 PA-X compared to CA07 PA-X in our assays, valine-100 is not the only marker for the increased PA-X activity in the nuclease domain. Based on our alignment presented in S1 Fig., BC24 PA-X N-terminal domain had highest identity to the PR8 N-terminal domain, with only 6 amino acid differences, all identical to CA07 PA-X. This makes it likely that the differences in the X-ORF between our H5N1-derived PA-X proteins and CA07 are responsible for increased shutoff activity and partially mitigate nuclease domain impairments due to baloxavir resistance mutations. In CA07 and all currently circulating seasonal H1N1 viruses, the X-ORF is shorter than in H3N2 or H5N1 strain PA-X proteins (41 vs. 61 amino acids). Given that the previous studies did not show significant effects of the X-ORF length on the PA-X shutoff activity [27], it is possible that the amino acid differences between the CA07 and BC24 X-ORFs are associated with different PA-X activity. By contrast, SW13 H3N2 PA-X had the highest identity to the PR8 among the analyzed PA-X proteins, with only a single substitution, alanine-20, present in CA07, TX24, and BC24 sequences. The remaining differences are unique to the SW13, indicating that the higher activity of H3N2 derived PA-X is caused by features that are distinct from those characterized for the PA-X proteins encoded by the avian-origin PA segments.

In our reporter assays, baloxavir resistance substitutions in TX24 and BC24 PA-X proteins significantly decreased their activity, but the magnitude of the remaining shutoff was higher than for the PR8 PA-X and comparable to the shutoff by the wild type 2009 pandemic H1N1 PA-X. This poses the question of whether baloxavir resistance mutations are associated with a lower fitness cost in H5N1 avian-origin strains and, if true, are they more likely to spread widely compared to the H1N1 viruses. So far, our data suggests that this may not be the case. First, our reporter shutoff assays rely on a single PA-X target and on ectopic overexpression of PA-X that is likely much higher than in virus-infected cells. Even under these conditions, we observed a sharp increase in 38T mutant PA-X expression levels compared to the WT, indicating much lower inhibition of its own expression. Second, we observed a clear impairment of host shutoff using the BC24 reconstituted minireplicon assay with 38T and 38M substitutions in PA. However, it is important that future studies, using IAVs possessing baloxavir resistance substitutions in PA derived from avian-origin H5N1 viruses, directly examine their relative fitness and the magnitude of host shutoff compared to the WT.

Our present study, as well as previous analyses, do not reveal significant defects in viral RdRp function caused by the baloxavir resistance mutations 38T and 38M, both present in the PA nuclease active site [18,19]. However, other analyses of the PA N-terminal domain endonuclease activity *in vitro* demonstrate that isoleucine 38 substitution mutants have much slower kinetics of substrate cleavage [12,19]. This catalytic activity is essential for the cap snatching function of the viral RdRp and is inhibited by baloxavir. Similarly, this catalytic activity is key to the PA-X host shutoff and, as shown in our study, unlike the RdRp function, 38T and 38M substitutions that affect the rate of nucleic acid cleavage significantly affect the PA-X activity. Why do these mutations affect PA-X function more than the RdRp function? We propose the following working model (Fig. 6E). The PA nuclease domain occupies a specific position on the surface of the trimeric RdRp complex and adopts optimal orientation for encountering RNA substrates [43]. These substrates, the nascent cellular capped pre-mRNAs, emerge from host RNA polymerase II, to which the viral RdRp is tethered through specific binding to the RNA polymerase II C-terminal domain [44]. Most importantly, the PB2 cap binding domain that is positioned next to the PA nuclease domain restricts lateral movement of the RNA by capturing the 5’ cap of the host pre-mRNA [43]. This may allow more time for the PA endonuclease active site to properly position the RNA substrate and catalyze RNA cleavage such that its catalytic rate is not the defining factor for overall cap snatching efficiency. By contrast, PA-X does not possess specific RNA binding domains and is recruited to its RNA targets by yet to be identified mechanisms that involve X-ORF interactions with host RNA processing factors [28]. The unstructured X-ORF is less likely to direct the RNA substrate towards the catalytic site effectively or restrict the lateral movement of the RNA for optimal cleavage. Therefore, the catalytic rate of the nuclease has major effects on the efficiency of cleavage of RNA targets that transiently interact with the active site. Additional support for this model comes from the analyses of the influenza B virus PA endonuclease domain activity which is ∼20 times slower than the influenza A virus endonuclease [19]. This does not prevent the influenza B virus RdRp from functioning similarly to the IAV RdRp. However, influenza B viruses do not encode PA-X [23]. It is possible that the requirement for maintaining PA-X activity applies stronger evolutionary pressure for maintaining higher catalytic activity to the IAV PA N-terminal domain compared to the influenza B virus PA.

Despite high rates of treatment-emergent baloxavir marboxil resistance due to PA active site mutations, sustained spread of resistant viruses have not been observed to date and this antiviral drug remains effective for treatment of influenza. Our study demonstrates that, while the effects of baloxavir resistance mutations on the viral RNA polymerase are negligible, they significantly impair activity of the viral host shutoff nuclease PA-X. Although the role of PA-X host shutoff in enabling sustained circulation and spread of IAV is not well defined [45,46], its function in supressing immune responses suggest that impaired host shutoff may represent an important barrier to the spread of baloxavir resistant IAVs.

Accordingly, for a widespread resistance to emerge, the compensatory mechanisms that augment PA-X function and/or potentiate other mechanisms of host shutoff and immune evasion must be present. Therefore, continued evaluation of the fitness of viral isolates with baloxavir resistance mutations should include assessment of viral host shutoff efficiency to assess their potential for spread in the population.

## MATERIAL AND METHODS

### Cell lines, viruses, and infections

The human lung adenocarcinoma A549 cells and the human embryonic kidney (HEK) 293A cells (American Type Culture Collection (ATCC), Manassas, VA, USA) were maintained in Dulbecco’s modified Eagle’s medium (DMEM), supplemented with heat-inactivated 10% fetal bovine serum (FBS) and 2 mM L-glutamine (all purchased from Thermo Fisher Scientific, Waltham, MA, USA) at 37°C in 5% CO2 atmosphere. Wild-type (WT) influenza virus A/PuertoRico/8/34(H1N1) (PR8) and the mutant viruses PR8-PA(fs), PR8-PA(38T), and PR8-PA(38M) were generated using an 8-plasmid reverse genetic system [47], as previously described in [24]. Individual segments of the A/California/07/2009(H1N1) (CA07) virus provided by the Public Health Agency of Canada (PHAC) National Microbiology Laboratory were subcloned into an 8-plasmid reverse genetic system. The WT CA07 and the mutant viruses CA07-PA(38T) and CA07-PA(38M) were generated from the reverse genetic system as described in [48]. In both PR8 and CA07 strain backgrounds, isoleucine 38 to threonine (38T) or methionine (38M) substitutions were introduced using PCR mutagenesis. A single nucleotide in the ATA codon was replaced to produce either the ACA (threonine) or the ATG (methionine) codon in plasmids encoding each of the PA segments. Virus stocks were produced in Madin Darby Canine Kidney (MDCK) cells (ATCC). Titers of all viral stocks were determined by plaque assays in MDCK cells using Avicel overlays as described in [49]. For each infection experiment, A549 cell monolayers in 12-well cluster dishes with or without glass coverslips were inoculated at an MOI of 1 for 1 h at 37°C. Then cells were washed briefly with phosphate-buffered saline (PBS, Thermo Fisher Scientific, Waltham, MA, USA) and cultured in infection medium (0.5% bovine serum albumin (BSA, Sigma-Aldrich, Missouri, USA) in DMEM) at 37°C, 5% CO2 atmosphere for 21 h prior to analysis.

### Plasmids

Expression plasmids encoding the C-terminally myc-tagged PA (pCR3.1-PA-myc) and PA-X (pCR3.1-PAX-myc) proteins from the PR8 strain and the pCR3.1-EGFP reporter were described previously [24,26]. Expression plasmids for the c-terminally tagged PA-X proteins from CA07, A/Switzerland/9715293/13(H3N2) (SW13), A/Bovine/texas/24-029328-01/24(H5N1) (TX24), and A/BC/PHL-2032/24(H5N1) (BC24) were generated by replacing the PR8 PA open reading frame in the pCR3.1-PAX-myc vector with synthetic gene fragments corresponding to PA open reading frames of each strain using KpnI and MluI restriction sites. Similarly, the C-terminally myc-tagged BC24 PA expression construct was generated by replacing the PR8 PA open reading frame in pCR3.1-PA-myc plasmid with synthetic BC24 PA gene fragment. The untagged BC24 PB2, PB1, and NP expression vectors were produced using the same strategy using the XhoI restriction site (downstream of the myc-tag) instead of the MluI site. The pCMV-LUC2CP/intron/ARE vector expressing firefly luciferase is a kind gift from Gideon Dreyfuss (Addgene plasmid #62858 [50]). To generate the RNA polymerase I-driven influenza A virus EGFP and firefly luciferase minireplicon reporters, the reverse genetic vector for the CA07 segment 8 was used. First, the CMV promoter and the NS1/NEP coding sequences downstream of the 8-th codon were deleted from this vector using PCR mutagenesis to generate pHWP1-N8 vector.

Subsequently, the PCR-amplified EGFP coding sequence from the pCR3.1-EGFP and the firefly luciferase coding sequence from the pTRE2-Fluc vector [51] were inserted downstream of the first 8 NS1/NEP codons into the pHWP1-N8 vector to produce pHWP1-N8-EGFP and pHWP1-N8-FF-luc minireplicon vectors, respectively. Full sequences of mutagenesis primers and vectors featured in this work are available upon request.

### Reporter transfections

For EGFP reporter shutoff assays, HEK 293A cells were seeded into 20-mm wells of 12-well cluster dishes and transfected the next day with 500 ng DNA mixes/well containing 100 ng pCR3.1-EGFP reporter, 350 ng pUC19 filler DNA, 50 ng of either the PA-X expression or empty control plasmid, and 1.5 ug of linear polyethylenimine (Cedarlane, Burlington, ON, Canada) in 100 ul of Opti-MEM-I media (Thermo). At 20-21 h post-transfection, live cells were imaged in the GFP channel using EVOS XL Core Imaging System (Thermo). For each independent biological replicate, duplicate wells were transfected with the same reporter DNA mix, and at least 2 separate random fields of view were captured per well. Total green fluorescence signal was measured for each field of view using ImageJ Fiji software after automatic background subtraction [52]. The integrated density of all pixels in the total of at least four images per replicate were averaged and used as a raw reporter expression value in further analysis. At least 3 independent biological replicates were performed per condition and the normalized values for each replicate were displayed on bar graphs. For luciferase reporter shutoff assays, HEK 293A cells were seeded into 96-well cluster dishes and transfected the next day with 60 ng DNA mixes/well containing 12 ng firefly luciferase reporter, 42 ng pUC19 filler DNA, 6 ng of either the PA-X expression or empty control plasmid, and 0.18 ug of linear polyethylenimine (Cedarlane, Burlington, ON, Canada) in 20 ul of Opti-MEM-I media (Thermo). Cells were analysed 24 h post-transfection using CLARIOstar Plus imager (BMG Labtech, Cary, NC, USA) with Promega Luciferase Assay System (E1501, Promega, Madison, WI, USA) according to the manufacturer’s protocol. At least 3 independent biological replicates were performed per condition and the normalized values for each replicate were displayed on bar graphs.

### Minireplicon assays

Reconstituted PR8, CA07, and BC24 minireplicon assays were performed in HEK 293A cells. For luciferase replicon assays, cells seeded in 96-well cluster dishes were transfected the next day with 60 ng DNA mixes/well containing 12 ng each of the expression vectors for PB2, PB1, PA, and NP proteins and the pHWP1-N8-FF-luc plasmid and 0.18 ug of linear polyethylenimine (Cedarlane, Burlington, ON, Canada) in 20 ul of Opti-MEM-Imedia (Thermo). Where indicated, at 4 h post-transfection cells were treated with various concentrations of baloxavir acid (BXA) by replacement of transfection media with growth media with or without BXA. Cells were analysed 24 h post-transfection using CLARIOstar Plus imager (BMG Labtech) with Promega Luciferase Assay System according to the manufacturer’s protocol. At least 3 independent biological replicates were performed per condition and the normalized values for each replicate were displayed on the graphs. For BC24 EGFP minireplicon, cells seeded in 12-well cluster dishes with glass coverslips were transfected with 500 ng/well of DNA mixes containing equal amounts of expression vectors for PB2, PB1, PA, and NP proteins and the pHWP1-N8-EGFP plasmid and 1.5 ug of linear polyethylenimine (Cedarlane, Burlington, ON, Canada) in 100 ul of Opti-MEM-I media (Thermo). At 24 h post-transfection, cells were fixed and analysed using immunofluorescence microscopy.

### Immunofluorescence microscopy

Cell fixation and immunofluorescence staining were performed according to the procedure described in [53]. Briefly, cells grown on 18 mm round coverslips were fixed with 4% paraformaldehyde in PBS for 15 min at ambient temperature and permeabilized with cold methanol for 10 min. After 1 h blocking with 5% bovine serum albumin (BSA, BioShop, Burlington, ON, Canada) in PBS, staining was performed overnight at +4°C with antibodies to the following targets: NP (1:1,000, mouse, Santa Cruz, sc-101352), PABPC1 (1:1000, rabbit, Abcam, ab21060). Alexa Fluor (AF)-conjugated secondary antibodies used were as follows: donkey anti-mouse IgG AF488 (Invitrogen, A21202), donkey anti-rabbit IgG AF555 (Invitrogen, A31572. Cell nuclei were stained with Hoechst 33342 dye (Invitrogen, H3570).

Slides were mounted with ProLong Gold Antifade Mountant (Thermo Fisher) and imaged using a Zeiss AxioImager Z2 fluorescence microscope. Green, red, blue, and far-red channel colors were changed for image presentation in the color-blind safe palette without altering signal levels. Quantification of cells with nuclear PABPC1 was performed as described in [29] from at least three randomly selected fields of view, analyzing >100 cells per replicate.

### Western blotting

Whole-cell lysates were prepared by the direct lysis of PBS-washed cell monolayers with 1x Laemmli sample buffer (50 mM Tris-HCl pH 6.8, 10% glycerol, 2% sodium dodecyl sulphate (SDS), 100 mM DTT, 0.005% Bromophenol Blue). Lysates were immediately placed on ice, homogenized by passing through a 21-gauge needle, and stored at –20°C. Aliquots of lysates thawed on ice were incubated at +95°C for 3 min, cooled on ice, separated using denaturing polyacrylamide gel electrophoresis, transferred onto polyvinylidene fluoride (PVDF) membranes using Trans Blot Turbo Transfer System with RTA Transfer Packs (Bio-Rad Laboratories, Hercules, CA, USA) according to the manufacturer’s protocol, and analyzed by immunoblotting using antibody-specific protocols. Antibodies to the following targets were used: β-actin (1:2000; HRP-conjugated, mouse, Santa Cruz, sc-47778); GFP (1:1000; rabbit, Cell Signaling, #2956); influenza A virus (1:2,000; goat, Abcam, ab20841, detect NP and M1 proteins); myc-tag (1:1000; mouse, Cell Signaling, #2276); NP (1:1000; mouse, Santa Cruz, sc-101352); PA (1:1000; rabbit, GeneTex, GTX125932). For band visualization, HRP-conjugated horse anti-mouse IgG (Cell Signaling, #7076), goat anti-rabbit IgG (Cell Signaling, #7074), or mouse anti-goat IgG (Santa Cruz, sc-2354) was used with Clarity Western ECL Substrate on the ChemiDoc Touch Imaging System (Bio-Rad Laboratories).

### RT-qPCR

Total RNA was extracted using the RNeasy Plus (Qiagen, Hilden, Germany) kit protocol according to the manufacturer’s instructions. 250 ng of total RNA was used to synthesize cDNA using LunaScriptRT SuperMix (New England BioLabs Inc, Massachusetts, USA).

Quantitative PCR amplification was performed using PerfeCta SYBR Green PCR master mix (QuantaBio, Beverly, MA, USA) and specific primers listed below on a Cielo 3 QPCR unit (Azure Biosystems, California, USA). Primers used: 18S-Left: cgttcttagttggtggagcg, 18S-Right: ccggacatctaagggcatca, ACTB-Left: catccgcaaagacctgtacg, ACTB-Right: cctgcttgctgatccacatc; G6PD-Left: tgaggaccagatctaccgca, G6PD-Right: aaggtgaggataacg caggc; IFNB1-left: gcctcaaggacaggatgaac; and IFNB1-right: agccaggaggttctcaacaa. Relative target levels were determined using the ΔΔCt method using 18S as a normalizer.

## SUPPLEMENTARY FIGURE LEGENDS

**S1 Figure. PA-X protein alignment.** (A) Sources of PA-X sequences and their amino acid identity. (B) BLASTP alignment of PA-X proteins from influenza A virus strains shown in panel A. PR8, CA07, SW13, TX24, and BC24 PA-X proteins are featured in this study; WS22 (H1N1) and DC23 (H3N2) are the recommended vaccine strains for the 2024-2025 season representing currently circulating viruses and are included for comparison. Positions of the isoleucine 38 and aspartate 108 in the nuclease active site are highlighted in yellow and red, respectively. Phenylalanine 191 marking the frame shift site and the beginning of the X-ORF is highlighted in grey.

